# Investigation of Bile Salt Hydrolase Activity in Human Gut Bacteria Reveals Production of Conjugated Secondary Bile Acids

**DOI:** 10.1101/2025.01.16.633392

**Authors:** Lauren N. Lucas, Jillella Mallikarjun, Lea E. Cattaneo, Bhavana Gangwar, Qijun Zhang, Robert L. Kerby, David Stevenson, Federico E. Rey, Daniel Amador-Noguez

## Abstract

Through biochemical transformation of host-derived bile acids (BAs), gut bacteria mediate host-microbe crosstalk and sit at the interface of nutrition, the microbiome, and disease. BAs play a crucial role in human health by facilitating the absorption of dietary lipophilic nutrients, interacting with hormone receptors to regulate host physiology, and shaping gut microbiota composition through antimicrobial activity. Bile acid deconjugation by bacterial bile salt hydrolase (BSH) has long been recognized as the first necessary BA modification required before further transformations can occur. Here, we show that BSH activity is common among human gut bacterial isolates spanning seven major phyla. We observed variation in both the extent and the specificity of deconjugation of BAs among the tested taxa. Unexpectedly, we discovered that certain strains were capable of directly dehydrogenating conjugated BAs via hydroxysteroid dehydrogenases (HSD) to produce conjugated secondary BAs. These results challenge the prevailing notion that deconjugation is a prerequisite for further BA modifications and lay a foundation for new hypotheses regarding how bacteria act individually or in concert to diversify the BA pool and influence host physiology.

## Introduction

The human gut microbiota plays a pivotal role in health and disease by biochemically transforming host-derived bile acids (BAs). Two primary BAs, cholic acid (CA) and chenodeoxycholic acid (CDCA), are synthesized in the human liver from cholesterol and conjugated to glycine or taurine (Bourgin et al., 2021; Foley et al., 2019). These conjugated BAs are stored in the gallbladder and released into the intestines following a meal, where they reach millimolar concentrations. A significant portion of conjugated BAs are reabsorbed in the terminal ileum and recirculated to the liver through the portal vein in a process called enterohepatic circulation. The remaining BAs in the small and large intestines are subject to deconjugation and transformation by gut bacteria. The transformed BAs are then either passively reabsorbed across the intestinal wall or excreted in feces.

Bile salt hydrolases (BSH) are bacterial enzymes that catalyze deconjugation by cleaving the amide bond in conjugated BAs to release unconjugated BAs. BSH activity has been linked to both positive and negative health outcomes in both humans and mice (Bourgin et al., 2021). BSH-active bacteria are reported to combat hypercholesterolemia (M. L. Jones et al., 2012) and non-alcoholic fatty liver disease (Huang et al., 2020). Elevated levels of primary BAs resulting from high BSH activity have been shown to stimulate hepatic NKT cell accumulation and antitumor immunity in mice (Ma et al., 2018). Additionally, BSH activity has been associated with resistance to *Clostridioides difficile* infection (Foley et al., 2023). In other contexts, positive health outcomes have been associated with limited BSH activity. For example, BSH deficiency in mice has been linked to reduced weight gain on a high-fat diet and increased lipid utilization over carbohydrates for energy (Yao et al., 2018). Reduced BSH activity has also been associated with slower progression of colorectal cancer (Y. Liu et al., 2022; Sun et al., 2023). These findings suggest that inhibiting BSH could be a therapeutic strategy for metabolic diseases. However, limited BSH activity may also lead to adverse outcomes, as elevated levels of conjugated BAs have been associated with inflammatory bowel disease (IBD) (Ogilvie & Jones, 2012), Type 2 diabetes (Labbé et al., 2014), and cholangiocarcinoma (CCA), an often fatal cancer of the biliary tract (R. Liu et al., 2014). These varied outcomes highlight the need for a comprehensive, systematic understanding of BSH activity across diverse genera of human gut bacteria.

Deconjugation by BSH has long been considered a “gateway” reaction (B. V. Jones et al., 2008) that allows unconjugated primary BAs to be further transformed into secondary BAs. Subsequent transformations produce BAs such as deoxycholic acid (DCA) and lithocholic acid (LCA) through dehydroxylation at the C7 position by enzymes encoded by the *bai* operon. Other transformations that occur are the oxidation of hydroxyl groups by ɑ-HSDs that generate position specific -oxoBAs, and the subsequent epimerization by β-HSDs to produce β-oriented BAs, such as ursodeoxycholic acid (UDCA) and ursocholic acid (UCA) (Ridlon et al., 2006). Additionally, BAs can be transformed into microbially conjugated BAs (MCBAs) (Lucas et al., 2021; Quinn et al., 2020) through the recently identified transferase activity of BSH enzymes (D. V. Guzior et al., 2024; Rimal et al., 2024).

Structural transformations of BAs profoundly influence their physiological roles. Glycine- and taurine-conjugated BAs are more hydrophilic than their unconjugated counterparts, facilitating their removal from the gastrointestinal (GI) tract and uptake by the liver during systemic circulation (Hofmann & Hagey, 2014; Ridlon & Bajaj, 2015; Zhou & Hylemon, 2014). While circulating throughout the body, BAs also interact with organs and tissues beyond the GI tract, including the brain, oral cavity, vagina, skin, and the nasal cavity (Mohanty et al., 2024). Through interactions with nuclear and membrane receptors, such as farnesoid X receptor (FXR), pregnane X receptor (PXR), and Takeda G-protein coupled receptor 5 (TGR5) (Ridlon et al., 2016), BAs regulate gene expression to influence cholesterol and glucose homeostasis, energy metabolism, inflammation, and xenobiotic metabolism (Björkholm et al., 2009; Bourgin et al., 2021). Oxidized BAs have been implicated in promoting colorectal cancer (Dong et al., 2024) and β-HSDs, which have reduced hydrophobicity, are less toxic to gut bacteria and have modulated agonistic or antagonistic interactions with host receptors (Doden & Ridlon, 2021). Due to the many roles of BAs in human health, there is a need for a more comprehensive understanding of BA transformations and dynamics by the gut microbiota.

To investigate the role of BSH activity in the diversification of the BA pool, we screened 77 gut bacterial strains spanning seven major phyla for their ability to deconjugate, transform, and reconjugate human conjugated bile acids. Due to its recognition as the rate-limiting BA transformation, we evaluated the ways in which altered BSH activity impacted the development of the BA pool for selected strains in time-course monoculture and coculture experiments. Our systematic evaluations generate new knowledge regarding bacterial bile acid deconjugation and transformations and provide a foundation for developing testable hypotheses that define causal links between the microbiome, bile acid pool composition, and human health.

## Results

### Bile salt hydrolase activity is widespread

We evaluated the deconjugation ability of 77 human gut bacterial strains (Supp. Table 1), spanning seven different phyla and 41 genera, against a mixture of the five most prevalent human conjugated BAs: taurocholic acid (TCA), glycocholic acid (GCA), taurochenodeoxycholic acid (TCDCA), glycochenodeoxycholic acid (GCDCA), and taurodeoxycholic acid (TDCA). Each strain was cultured anaerobically in a medium supplemented with conjugated BA pools at 100 µM and 500 µM concentrations. Cultures were grown until they reached stationary phase, at which point samples were collected for LC-MS/MS analysis.

Bacterial species were considered to have BSH activity if they deconjugated more than 2% of the provided conjugated BAs. BA deconjugation activity was widespread, with BSH activity detected in 56 of the 77 tested bacterial strains, spanning the Actinomycetota, Bacillota, Bacteroidota, Fusobacteria, and Pseudomonadota phyla (Fig.1). A single species from the Lentisphaerota and Verrucomicrobiota phyla was tested, and neither exhibited BSH activity. We observed deconjugating activity on both glycine- and taurine-conjugated BAs, with some bacterial species showing a preference for one over the other. Patterns in deconjugating activity were similar across the 100 µM and 500 µM concentrations, with few exceptions. Beyond the production of unconjugated primary BAs through deconjugation, several species performed further transformations to the BA core to produce unconjugated secondary BAs (Fig. 1) and/or reconjugated BAs to generate microbially conjugated bile acids (MCBAs) (Fig. 2, Supp. Fig. 3). Surprisingly, we identified conjugated secondary BAs in several samples (Fig.1, Supp. Fig. 1, 2).

**Figure 1.**
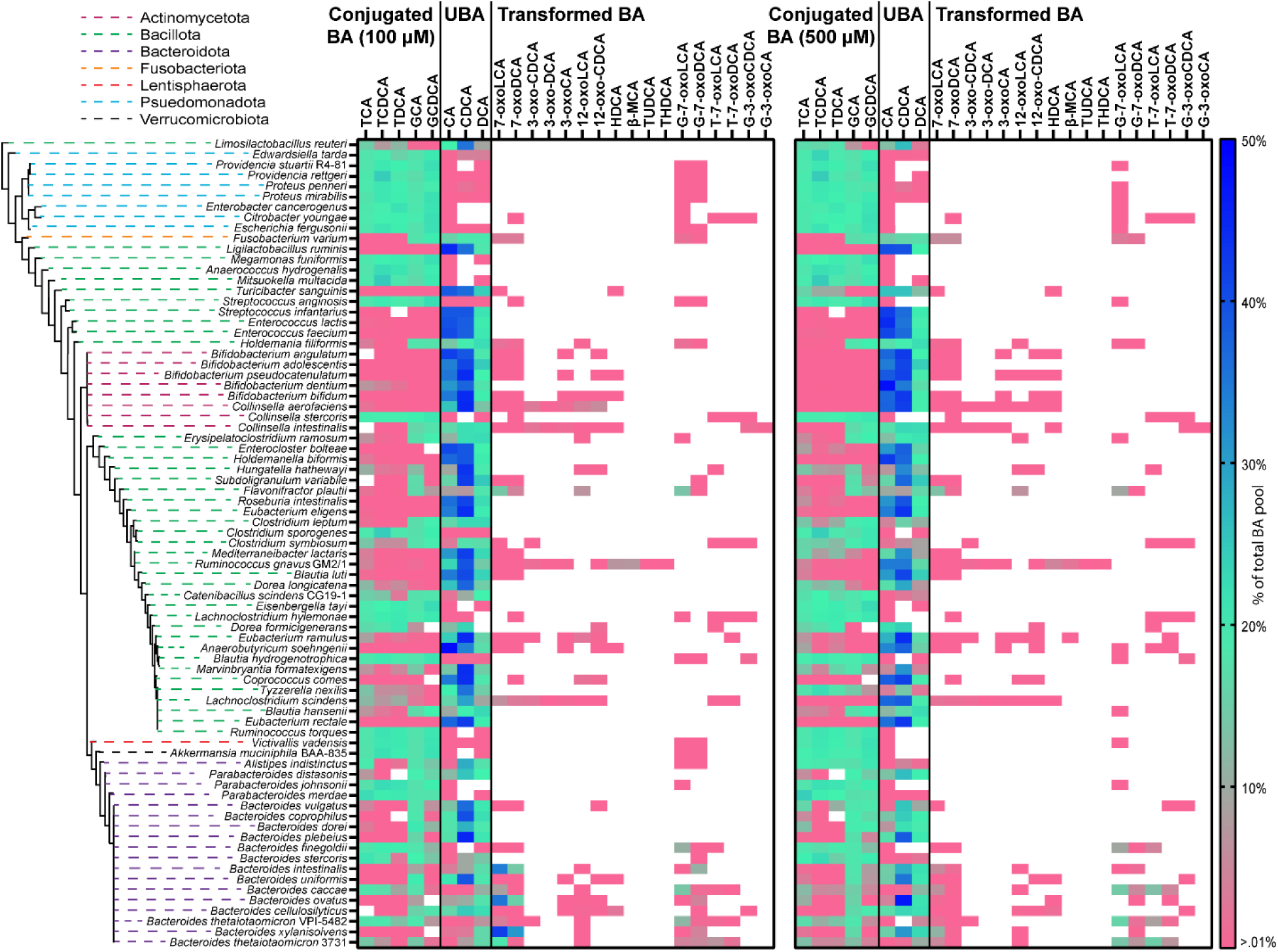
Bile acid deconjugation and transformation measurements across phyla. The heat maps show percent of bile acids (BAs) relative to total starting conjugated BA concentration, 100 µM (left) or 500 µM (right), as measured by LC-MS using external standard calibration curves. There were two replicates for each strain at each concentration. For each heatmap, conjugated BA substrate measurements are depicted in the left column, unconjugated BAs (UBA) in the middle column, and transformed BAs in the right column. Color scale denotes BAs up to 50% as dark blue, 20% in green, >.01% in pink, and below detection in white. Phyla information is indicated by color-coded dashed lines in the phylogenetic tree. Bile acid name abbreviations: TCA, taurocholic acid; TCDCA, taurochenodeoxycholic acid; TDCA, taurodeoxycholic acid; GCA, glycocholic acid; GCDCA, glycochenodeoxycholic acid; CA, cholic acid; CDCA, chenodeoxycholic acid; DCA, deoxycholic acid; 7-oxoLCA, 7-oxolithocholic acid; 7-oxoDCA, 7-oxodeoxycholic acid; 3-oxoCDCA, 3-oxochenodeoxycholic acid; 3-oxoDCA, 3-oxodeoxycholic acid; 3-oxoCA, 3-oxocholic acid; 12-oxoLCA, 12-oxolithocholic acid; 12-oxoCDCA, 12-oxochenodeoxycholic acid; HDCA, hyodeoxycholic acid; bMCA, beta-muricholic acid; TUDCA, tauroursodeoxycholic acid; THDCA, taurohyodeoxycholic acid; G-7-oxoLCA, glyco-7-oxolithocholic acid; G-7-oxoDCA, glyco-7-oxodeoxycholic acid; T-7-oxoLCA, tauro-7oxolithocholic acid; T-7-oxoDCA, 7-oxodeoxycholic acid; G-3-oxoCDCA, glyco-3-oxochenodeoxycholic acid; G-3-oxoCA, glyco-3-oxocholic acid. See **Supplementary Figure 1** for bile acid transformations and structures.

**Figure 2.**
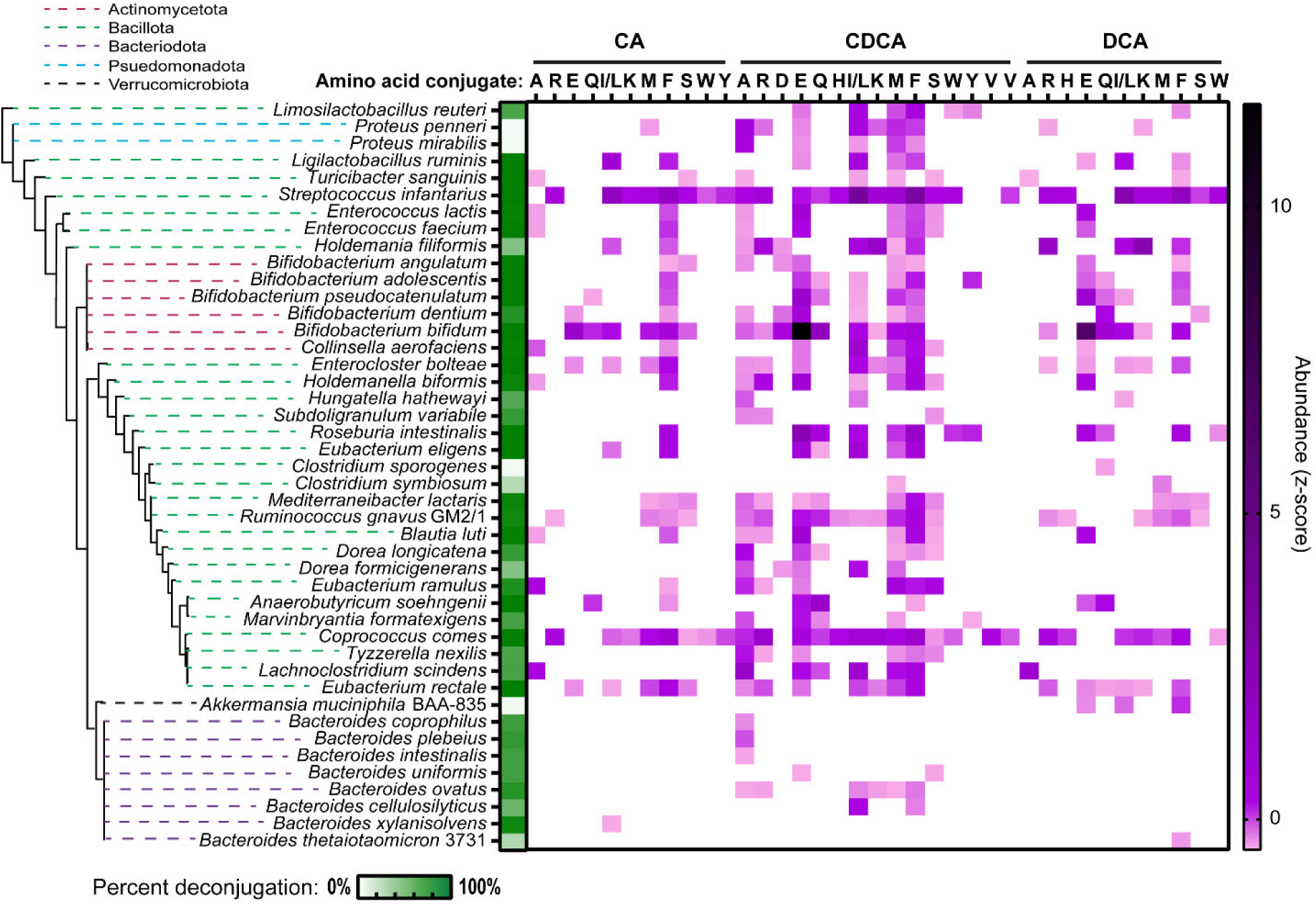
Microbially conjugated bile acid production at 100 µM. The heat map shows relative levels of MCBAs produced, as denoted by z-score, with the mean and standard deviation calculated from raw signal across samples. Phyla information is indicated by color-coded dashed lines in the phylogenetic tree. Amino acid conjugations are indicated by their single letter abbreviation across the top of the heat map. Glycine and taurine conjugations could not be measured in this study because these conjugated BAs were provided in the media. The heat map for BAs at 500 µM can be found in **Supplementary Figure 3**.

Of the 53 species previously identified to possess a putative *bsh* gene based on computational analyses (Heinken et al., 2019; Song et al., 2019), 44 exhibited BSH activity in our *in vitro* analysis (Supp. Table 2). Among the 24 species without a previously identified *bsh* gene, 12 exhibited activity. The highest levels of deconjugation were observed in the Actinomycetota, with more variable activity in the Bacillota and the Bacteroidota (Fig. 1). All *Bifidobacteria* and two out of three *Collinsella* species tested were able to deconjugate a majority of provided conjugated BAs to produce unconjugated BAs. Among the Bacillota, *Enterococcus* and *Eubacterium* species effectively deconjugated all provided conjugated BAs, as did *Enterocloster bolteae*, *Roseburia intestinalis*, *Anaerobutyricum soehngenii*, and *Coprococcus comes*, consistent with prior studies (Majait et al., 2023; D. Wang et al., 2021). The probiotic genus *Lactobacilli* also exhibited BSH activity, with *Lactobacillus ruminis* deconjugating all conjugated BAs, while *Lactobacillus reuteri* showed a preference for glycine-conjugated BAs. In the Bacteroidota, 75% of species demonstrated BSH activity, deconjugating measurable amounts of unconjugated BAs (Fig. 1).

We observed limited BSH activity in the Pseudomonadota. *Edswardsiella tarda* deconjugated ∼16% of conjugated BAs when provided 100 µM conjugated BAs, but less than 2% when provided 500 µM. Similarly, *Proteus penneri* deconjugated only ∼5% of unconjugated BAs at both concentrations (Fig. 1, Supp. Table 2). *Fusobacterium varium,* our only representative from the Fusobacteriota phylum, exhibited deconjugating activity only on taurine-conjugated BAs. The majority of *Bifidobacterium* and *Enterococcus* species are known to possess a *bsh* gene, while *Bacteroides*, *Collinsella*, *Lactobacilli*, and *Streptococcus* are known to have genus level variation in BSH activity (Franz et al., 2001; Heinken et al., 2019; Kingkaew et al., 2023; Knarreborg et al., 2002; Li et al., 2021; Patterson et al., 2022; Ridlon et al., 2020; Ruiz et al., 2021; Shimada et al., 1969; Song et al., 2019; Wegner et al., 2017; Wijaya et al., 2004).

Unconjugated secondary BAs were produced through a combination of deconjugation and subsequent transformations (Fig. 1). The Actinomycetota, Bacillota, and Bacteroidota exhibited both the highest levels of deconjugation and highest production of secondary BAs. Notably, -oxoBA production was more prevalent at 100 µM than at 500 µM (Fig. 1). *Collinsella aerofaciens, Flavonifractor plautii, Lachnoclostridium scindens, Bacteroides intestinalis, Bacteroides ovatus,* and *Bacteroides xylanisolvens* showed the highest levels of secondary BA production. However, many species, including *Collinsella intestinalis*, *Holdemania filiformis*, *Dorea formicigenerans*, and *Erysipelatoclostridium ramosum*, exhibited high BSH activity but limited secondary BA production. While *B. finegoldii* and *B. hydrogenotrophica* produced conjugated secondary BAs, *Ruminoccocus torques, Lachnoclostridium hylemonae, Collinsella stercoris,* and *Escherichia fergusonii* did not generate expected secondary BAs, likely due to insufficient unconjugated BA substrates. For these species, BSH activity appeared to be the rate-limiting step for further transformations to occur (Batta et al., 1990; Begley et al., 2006; Foley et al., 2019).

Unexpectedly, we identified conjugated 7-oxo and 3-oxoBAs in our samples (Fig. 1, Supp. Fig. 1, 2). At 100 µM, 12 species produced conjugated -oxoBAs exceeding 1% of the provided conjugated BAs, while 9 species did so at 500 µM (Fig. 1, Supp. Table 2). Production of these BAs could result from either 1) deconjugation, transformation, and subsequent reconjugation of BAs, or 2) direct dehydrogenation of the C3 and C7 hydroxyl groups on conjugated BAs by hydroxysteroid dehydrogenases (HSDs). Notably, *Bacteroides finegoldii,* and to a lesser extent *Blautia hydrogenotrophica,* produced conjugated secondary BAs without detectable BSH activity, suggesting that HSDs can act directly on conjugated BAs. This observation challenges the widely accepted view that deconjugation is a prerequisite for further BA modifications. The remaining species exhibited both BSH activity and HSD activity, preventing us from determining, based on a single time point of data, whether HSD activity occurred on conjugated BAs, unconjugated BAs, or both (Fig. 1).

### Deconjugation correlates with MCBA production

In recent years, a new mechanism by which bacteria diversify the BA pool was discovered: the conjugation of BAs to amino acids to produce microbially conjugated BAs (MCBAs), (Lucas et al., 2021; Quinn et al., 2020). More recently, BSHs were identified as the enzymes responsible for this conjugation or transferase activity (Foley et al., 2023; D. Guzior et al., 2022; Patterson et al., 2022). Our analysis revealed that 45 out of the 77 bacterial species assessed for BSH activity were capable of conjugating a wide range of amino acids to CA, CDCA, and DCA (Fig. 2, Supp. Table 2). In general, species with high deconjugating activity also exhibited high conjugating activity, resulting in MCBA production. Similar to other BA transformations, a greater diversity of MCBAs was produced at 100 µM than at 500 µM (Fig.2, Supp. Fig. 3). Across both concentrations, amino acids were most frequently conjugated to CDCA, followed by DCA and then CA. Phenylalanine, alanine, glutamate, leucine/isoleucine, and methionine were the most frequently conjugated amino acids. The Bacillota and Actinomycetota phyla produced the highest levels and most diverse MCBAs, while the Pseudomonadota and Bacteroidota produced fewer and less diverse MCBAs (Fig. 2). Notably, *Streptococcus infantarius, Bifidobacterium bifidum,* and *C. comes* produced the highest concentrations and most diverse MCBAs.

There were some exceptions to the correlation between deconjugation activity and MCBA production. Specifically, seven members of Bacteroidota, five of Bacillota, one of Actinomycetota, and one of Fusobacteroidota deconjugated BAs but did not produce MCBAs (Supp. Table 2). In contrast, some species with bioinformatically identified *bsh* did not exhibit deconjugating activity but still produced MCBAs, presumably through the enzyme’s transferase activity. These species were *Providencia rettgeri*, *Proteus mirabilis*, *Clostridium sporogenes*, and *Akkermansia muciniphila*. Of these, only *P. mirabilis* exhibited MCBA production at 500 µM (Supp. Fig. 3). Other species, including *Bacteroides coprophilus*, *Bacteroides plebeius*, *B. intestinalis*, *Bacteroides thetaiotaomicron* 3731, *Bacteroides adolescentis*, and *Clostridium symbiosum* lost MCBA production at 500 µM (Supp. Fig. 3). Finally, several species with newly discovered deconjugating activity also produced MCBAs, including *H. filiformis, Subdoligranulum variabile, C. symbiosum, Blautia luti, Eubacterium ramulus, Bacteroides cellulosilyticus*, and *B. thetaiotaomicron* 3731.

### Deconjugation specificity follows phylogenetic patterns

Taurine-conjugated and glycine-conjugated BAs have variable levels of toxicity, and through BSH deconjugation specificity bacteria have been shown to competitively colonize the intestines (Foley et al., 2021, 2022; Grill et al., 2000). Moreover, BA conjugation type modulates interactions with host receptors that impact host physiology (Ridlon et al., 2016; H. Wang et al., 1999). We determined the BSH deconjugation specificity preferences toward taurine- and glycine-conjugated BAs for all BAs combined, and for each BA core, CDCA or CA (Fig. 3). Species were considered to have BSH specificity if they deconjugated 10% more of one type over the other. At the 100 µM concentration, 26 species showed a preference for taurine-conjugated BAs, while 8 preferred glycine-conjugated BAs. At 500 µM, 22 species preferred taurine-conjugated BAs and 11 preferred glycine-conjugated BAs. In addition, on average, 50% more taurine-conjugated BAs were deconjugated than glycine-conjugated BAs. This pattern was consistent across both BA cores (e.g., CDCA vs. CA).

**Figure 3.**
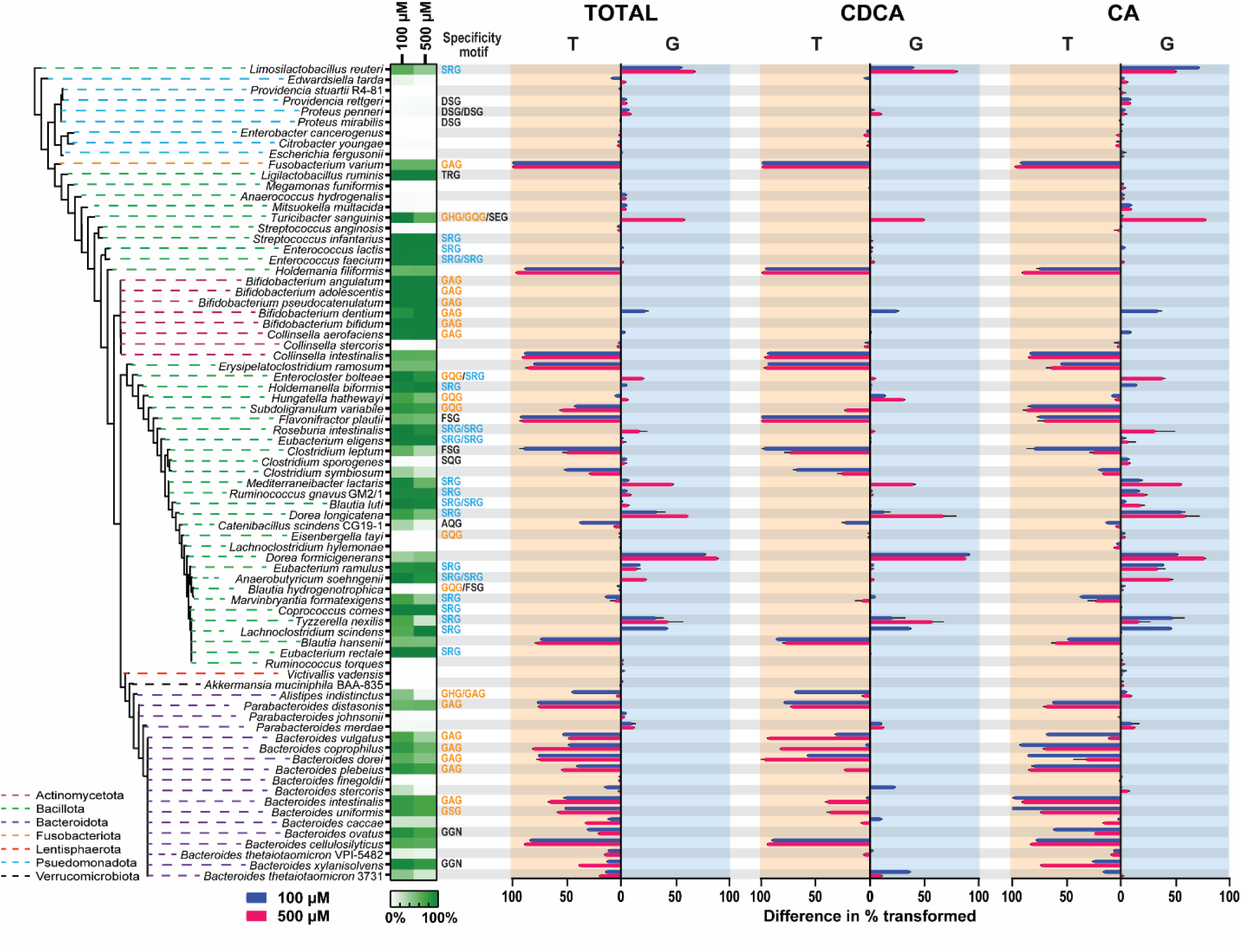
Bacterial specificity for glycine- and taurine-conjugated bile acids across phyla. Percent of glycine deconjugation was subtracted from taurine deconjugation for each strain at each concentration and the absolute value was plotted to show BSH preference (left column). This process was repeated for conjugated BAs with a CDCA core (middle column) and with a CA core (right column). A preference for taurine conjugated BAs is indicated by an orange shaded background, while a preference for glycine-conjugated BAs is shaded in blue, each corresponding with the predicted specificity motif of the same color. Values are averaged across two replicates and range bars are included. Values for the 100 μM condition are in blue and for the 500 μM condition in pink. Strains are organized by phylogeny as indicated by the colored dashed lines of the tree. The heat map indicates percent of BAs deconjugated for each strain, regardless of specificity.

Phylogenetic patterns in deconjugation specificity were apparent in the Bacteroidota and Bacillota. Of the 16 *Bacteroides* species that exhibited deconjugation specificity, only one showed a preference for glycine-conjugated BAs. Across both concentrations, *Bacteroides* species deconjugated ∼47% more taurine-conjugated BAs than glycine-conjugated BAs, consistent with previous reports (Yao et al., 2018). In contrast, deconjugation specificity within the Bacillota phylum was split, with a similar number of species showing preference for either taurine- or glycine-conjugated BAs. On average these species deconjugated approximately 25% more taurine-conjugated BAs at 100 µM and 12% more at 500 µM.

In general, bacteria exhibited more complete deconjugation at 100 µM than at 500 µM (Fig. 3). This pattern applied to species in the Bacillota (*Clostridium leptum*, *C. symbiosum*, *Mediterraneibacter lactaris*, *Dorea longicatena*, *Catenibacillus scindens* CG19-1, and *Tyzzerella nexilis*), the Bacteroidota (*Alistipes indistinctus*, *Bacteroides vulgatus*, and *Bacteroides stercoris*), and the Pseudomonodota phyla (*E. tarda*). Interestingly, at the 500 µM concentration, many species exhibited increased deconjugation specificity (Fig. 3). For example, *E. bolteae*, *Turicibacter sanguinis*, and *M. lactaris* were able to deconjugate all BAs at 100 µM, but exhibited glycine preference at 500 µM. In addition, at the 500 µM concentration, Bacteroidota species were more likely to deconjugate TCDCA than GCDCA, while at 100 µM conjugation preference was less pronounced. Conversely, and counterintuitively, *Bifidobacterium dentium* and *L. scindens* exhibited increased glycine specificity at 100 µM, yet complete deconjugation, and thus no specificity, at the 500 µM concentration. Altogether, these observations suggest a role of environmental conditions in BSH activity, with differences in concentrations of BAs leading to different levels of activity, potentially as a result of enzyme induction (Lundeen & Savage, 1990) or saturation (Stellwag & Hylemon, 1976).

We performed an *in silico* analysis to compare predicted and observed deconjugation specificity. For this, we compiled the genomes of all 77 strains in this study and identified their BSH protein sequences from the RefSeq NCBI database using the keywords “bile salt hydrolase” and “choloylglycine hydrolase”. We then built a hidden Markov model (HMM) using 84 BSHs from a previous study that found taurine-preferring BSH to have the motif “G-X-G” with X=T/V and glycine-preferring enzymes to have “S-R-X” with X=G/S (Foley et al., 2022). Our analysis identified one BSH in 39 species, two BSHs in nine species, and three BSHs in one species (Fig. 3). The BSHs in our species were predicted as taurine-preferring if they contained the motif “G-X-G” (with X=A/H/Q/S) and glycine-preferring if they contained the motif “S-R-G”. Among the identified BSHs, 23 were taurine preferring, spanning four phyla, and 24 were glycine preferring, all in the Bacillota phylum.

For most phyla, there was agreement between the predicted and observed BSH deconjugation specificity (Fig. 3). In the Bacteroidota and Fusobacteriota, the identified BSHs were predicted to have taurine specificity, which was consistent with our observed *in vitro* activity. In the Lentishpaerota, and Verrucomicrobiota, no *bsh* were identified, and no deconjugation activity was observed. In several members of the Bacillota, although conjugation preference varied, it was accurately predicted. For example, the BSH from *S. variabile* contained the G-Q-G motif, associated with taurine specificity, while the BSH from *D. longicatena* had the S-R-G motif, associated with glycine specificity, both of which were consistent with the observed activity. However, *T. sanguinis* was predicted to have taurine preference in two out of its three BSHs, yet only exhibited glycine preference at 500 µM.

Discrepancies between predicted and observed activity were most notable when bacteria were able to fully deconjugate all BAs, such as those in the Actinomycetota and in several species in the Bacillota. Most species in the Actinomycetota were broadly predicted to have taurine specificity, but deconjugated all supplied BAs. Similarly, in the Bacillota, *Entercoccus* species, along with several others, were predicted to have glycine specificity but deconjugated all BAs, while *Eisenbergella tayi* and *B. hydrogenotrophica* were predicted to have taurine specificity but did not deconjugate BAs (Fig. 3). The D-S-G motif in BSHs from the Pseudomonadota did not appear to confer deconjugating activity. Conversely, in some species a *bsh* was not identified, yet deconjugation specificity was still observed. For example, *D. formicigenerans* did not have an identified *bsh*, but showed strong glycine preference, while *C. intestinalis, H. filiformis*, *E. ramosum*, and *Blautia hansenii* did not have an identified *bsh*, but exhibited strong taurine preference. In addition, the motif F-S-G may also confer taurine specificity, as seen with *F. plautii* and *C. leptum*. Interestingly, BSHs with the F-S-G motif clustered more closely with glycine-preferring BSHs in other Bacillota species. These findings provide further support for the association of specific motifs with deconjugation preferences across phyla.

### Investigation of BSH dynamics in monoculture

Previous studies indicate that BSHs are intracellular enzymes with activity typically coupled to growth (Begley et al., 2005). BSH activity is recognized as a necessary action preceding further BA transformations (Batta et al., 1990; Foley et al., 2019; B. V. Jones et al., 2008). To investigate dynamics of BSH activity in relation to bacterial growth, we selected seven phylogenetically diverse species and monitored them over a period of 72 hours. Each species was cultured with five conjugated BAs, each at a concentration of 100 µM. Cultures were sampled to quantify BAs and measure optical density. These experiments revealed nuanced deconjugation dynamics across species and identified two distinct patterns of BSH-mediated secondary BA production (Fig. 4).

**Figure 4.**
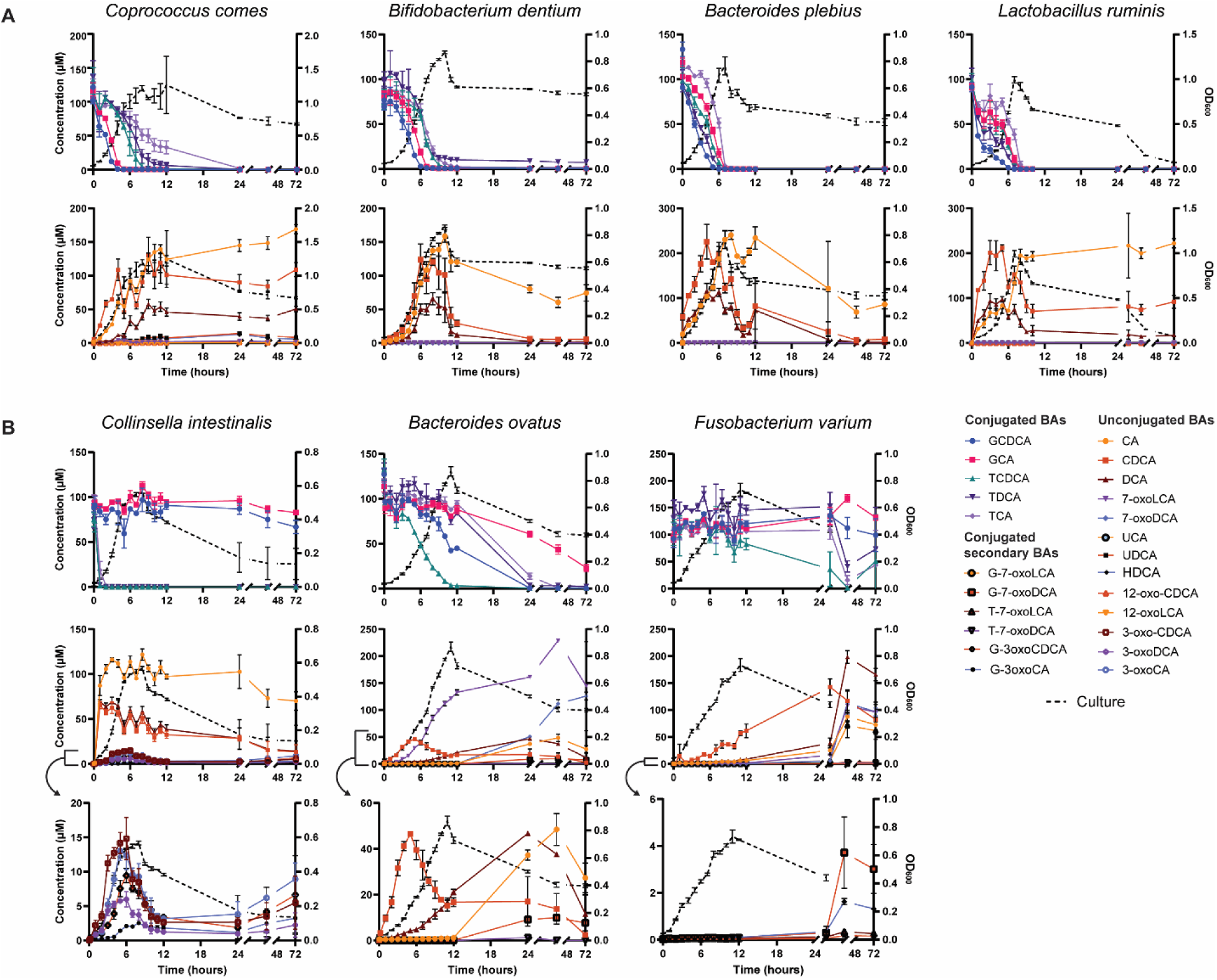
BSH dynamics in monoculture. A) Bile acid transforming activity for four species with complete BSH activity. B) Bile acid transforming activity for three species with incomplete, or specific, BSH activity. Low level transformations are shown in the bottom row. Strains with differing BSH activity were grown in triplicate and sampled over the course of 72 hours. Error bars represent the standard deviation of each averaged measurement. Figure legend shows which BA measurements are presented in each graph. Culture growth as measured by optical density is indicated by a black dashed line. Conjugated BAs in the top row and transformed BAs in the bottom row(s).

#### Timing of BSH activity varies across taxa

While four species -*C. comes*, *B. dentium*, *B. plebeius*, and *L. ruminis*-exhibited BSH activity coupled with growth and deconjugated all 5 conjugated BAs during exponential phase (Fig. 4A), the other three species -*C. intestinalis*, *B. ovatus*, and *F. varium*-did not follow the same pattern (Fig. 4B). Among species that deconjugated all 5 BAs, substrate preferences could be discerned. For instance, *C. comes* preferred to deconjugate glycine-conjugated BAs over taurine-conjugated BAs, with a ∼3-hour lag between the deconjugation of each type. *B. dentium* showed the same preference, but with a shorter gap of ∼1.5 hours between glycine- and taurine-conjugated BAs. *B. plebeius* and *L. ruminis* showed less distinct preferences, but appeared to deconjugate GCDCA first and TCA last, similar to *C. comes* (Fig. 4A). These data highlight the importance of time-series analyses in determining enzyme specificity, especially when deconjugation goes to completion.

The remaining three species had varied BSH patterns, either not coupling deconjugation to growth, or failing to deconjugate all five BAs within 72 hrs (Fig. 4B). *C. intestinalis* coupled BSH activity to growth but displayed a marked substrate preference, rapidly deconjugating all taurine-conjugated BAs while leaving glycine-conjugated BAs untouched. *B. ovatus* deconjugated glycine- and taurine-conjugated CDCAs during exponential growth, but did not deconjugate the remaining BAs until it reached stationary phase at 11 hours. Finally, *F. varium* deconjugated approximately half of the provided TCDCA during exponential growth and the remainder during stationary phase. Between 24 and 72 hrs, it also deconjugated the other taurine-conjugated BAs, but it is unclear if glycine-CBAs were deconjugated, as their disappearance may be explained by HSD activity to produce conjugated secondary BAs. These findings highlight that while many bacteria couple BSH activity to growth, others do not. The timing and specificity of BSH activity has significant implications for the network of BA transformations by gut bacteria in *in vivo* systems.

#### BSH activity correlates with MCBA production

BSHs have been identified as the enzymes responsible for conjugating BAs to amino acids to produce microbially conjugated BAs (MCBAs) (D. V. Guzior et al., 2024; Rimal et al., 2024). Our data show that levels of BSH activity directly correlated with MCBA production. The four species exhibiting rapid BSH activity and complete deconjugation of all BAs produced MCBAs, while the three species with slower or incomplete BSH activity did not (Fig. 4, Supp. Fig. 4). Estimated MCBA concentrations reached 8 µM for *C. comes*, 0.4 µM for *B. dentium*, 1.4 µM for *B. plebeius*, and 6 µM for *L. ruminis*, with production peaking concurrently with the levels of unconjugated BAs (Fig. 4A). Consistent with previous studies, the most abundant MCBAs were conjugated to the amino acids glutamate, glutamine, alanine, and asparagine (D. V. Guzior et al., 2024). Out of the seven species we analyzed, those with highest BSH activity lacked other types of BA transforming activity. This observation suggests that the combination of robust BSH activity, coupled with the availability of unconjugated BAs that are not diverted toward secondary BA production, promoted MCBA formation.

#### HSD activity produces conjugated-oxoBAs

While HSD activity is known to produce -oxoBAs from unconjugated BAs, we were surprised to observe that species with slow-acting or incomplete BSH activity also generated conjugated-oxoBAs throughout the time-course. The presence of slow-acting or incomplete BSH activity resulted in a mixed pool of unconjugated and conjugated BAs, enabling *C. intestinalis*, *B. ovatus*, and *F. varium* to simultaneously produce both conjugated and unconjugated-oxoBAs via HSD activity (Fig. 4B).

*C. intestinalis* rapidly deconjugated taurine-conjugated BAs, but exhibited no activity against glycine-conjugated BAs. As expected, CDCA, DCA, and CA were transformed into 3-oxoCDCA, 3-oxoDCA, and 3-oxoCA, respectively. Concurrent with the production of unconjugated -oxoBA production, G-3-oxoBAs, G-3-oxoCDCA and G-3-oxoCA, appeared in the media (Fig. 4B). Their presence indicated that the *C. intestinalis* 3ɑ-HSD was active on both conjugated and unconjugated BAs present in the media. The presence of glycine-conjugated secondary BAs is unlikely to result from reconjugation of 3-oxoBAs, as *C. intestinalis* BSH activity did not deconjugate glycine-conjugated BAs or produce MCBAs (Fig. 2), suggesting that its BSH activity is specific to taurine. Interestingly, all -oxoBAs were transient, peaking at ∼15 µM at 6 hrs but dropping below 5 µM by 12 hrs of growth. Concentrations increased again after 24 hrs, suggesting that *C. intestinalis* HSD activity may be reactivated during stationary phase.

Similarly, BSH activity in *B. ovatus* and *F. varium* produced a mixed pool of unconjugated and conjugated BAs (Fig. 4B). *B. ovatus* BSH deconjugated BAs in a staggered manner, preferring CDCAs, then CAs and DCA. CDCA accumulated in the media before it was transformed into 7-oxoLCA via 7ɑ-HSD activity. Subsequently, DCA and then CA levels increased, and after 12 hrs, both 7-oxoDCA and G-7-oxoDCA were produced. The concurrent production of conjugated and unconjugated-oxoBAs suggests that the *B. ovatus* HSD was active on both conjugated and unconjugated BAs. While reconjugation could explain the production of G-7-oxoDCA, it is unlikely because G-7-oxoLCA was not observed. *F. varium* followed a similar pattern to *B. ovatus*, preferentially deconjugating TCDCA to release CDCA throughout its growth. In addition, *F. varium* HSD activity simultaneously produced 7-oxoBAs from CA and CDCA, as well as G-7-oxoLCA and G-7-oxoDCA from GCDCA and GCA, respectively (Fig. 4B). The modest increase in conjugated BAs between 48 and 72 hrs could be attributed to reconjugation activity, but could also be explained by reversible HSD activity, reforming conjugated primary BAs from conjugated secondary BAs. Either mechanism broadens the known repertoire of BA transformations performed by gut bacteria.

### Coculture experiments reveal BSH impact on BA pool

Although it is widely accepted that bacteria with distinct BA transforming capabilities perform sequential modifications on BAs, direct experimental evidence for this process remains limited (Heinken et al., 2019; MacDonald et al., 1982; Ridlon et al., 2006). To directly examine whether sequential transformations take place and assess the impact of BSH activity on secondary BA production, we cocultured bacteria with varying levels of BSH activity — *Bifidobacterium angulatum*, *C. aerofaciens, S. infantarius*, and *C. symbiosum*— alongside *B. thetaiotaomicron* VPI-5482 (*B. theta*), which exhibits limited BSH activity but robust secondary BA production via 7ɑ-HSD activity. Each species was provided with 100 µM of each of the five conjugated BAs and cultured individually and in coculture for 72 hours. Cell growth was monitored by optical density, and samples were collected at regular intervals for LC-MS/MS analysis of BA concentrations.

The levels and timing of BSH activity directly influenced secondary BA production. When cultured alone, *B. angulatum* fully deconjugated all provided conjugated BAs within ten hours, generating CA, CDCA, and DCA (Fig. 5A). In coculture with *B. theta*, starting at approximately three hours of growth, CA and CDCA were transformed into 7-oxoDCA and 7-oxoLCA, respectively, (Fig. 5C). Together, these two species were able to transform conjugated primary BAs into unconjugated secondary BAs, a process that neither species could achieve independently.

**Figure 5.**
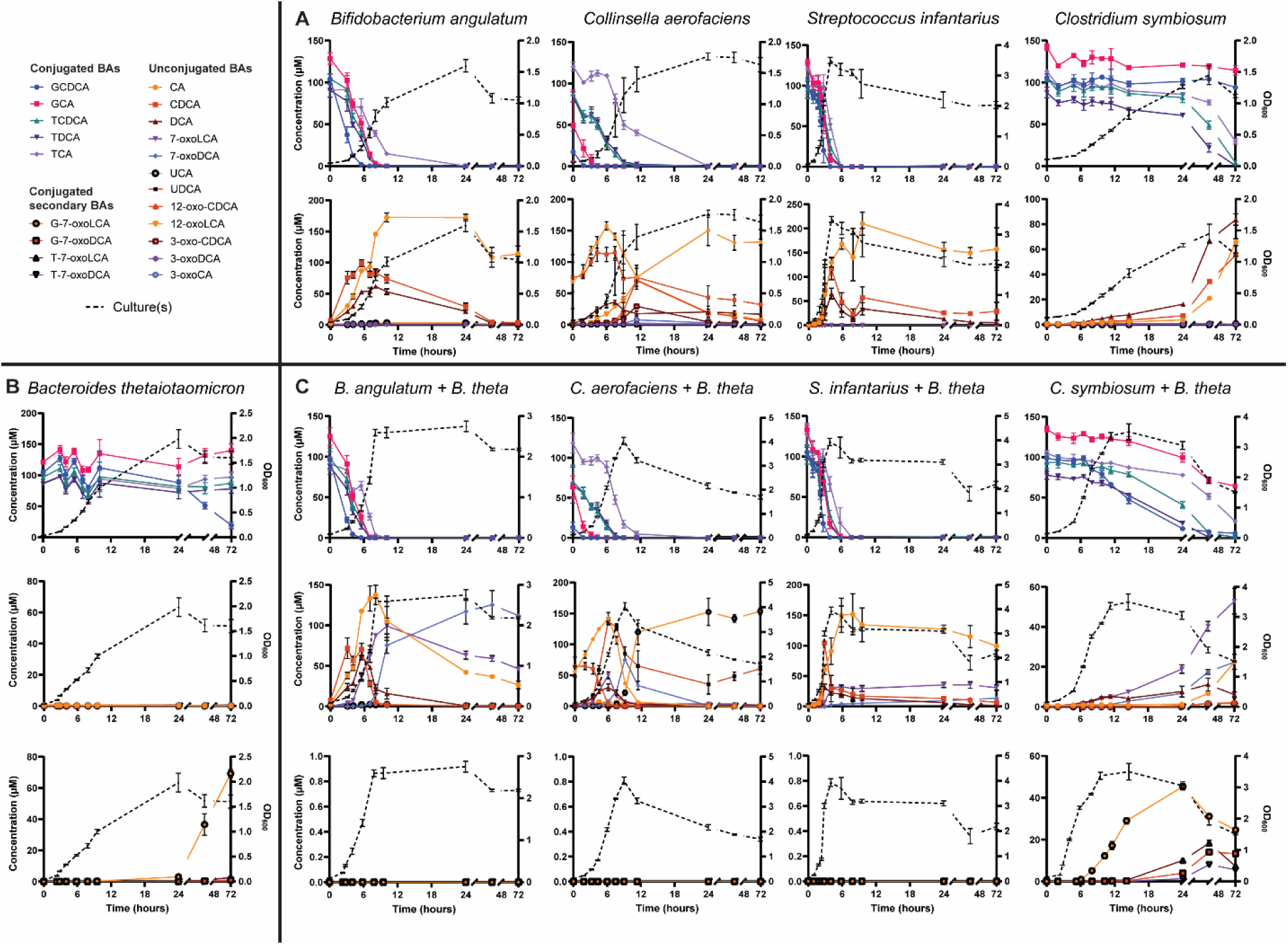
BSH dynamics in coculture. A) Bile acid transforming activity throughout growth for the four species with BSH activity. B) Bile acid transforming activity for *B. thetaiotaomicron*, which has HSD activity. C) Bile acid transforming activity for each coculture. Strains with differing BA transforming capabilities were grown in triplicate individually and in coculture and sampled over the course of 72 hours. Error bars represent the standard deviation of each averaged measurement. The legend lists bile acids grouped by how they are graphed. Culture growth as measured by optical density is indicated by a black dashed line.

*C. aerofaciens* deconjugated all conjugated BAs to produce unconjugated BAs and exhibited faster rates of deconjugation for glycine-conjugated BAs compared to taurine-conjugated BAs in both monoculture and coculture (GCDCA > GCA > TCDCA = TDCA > TCA) (Fig. 5). In monoculture, *C. aerofaciens* also converted unconjugated BAs into 12-oxoBAs and 3-oxoBAs (Fig. 5A). However, in coculture with *B. theta*, CA and CDCA were transformed into 7-oxoDCA and 7-oxoLCA by *B. theta*, which were then epimerized by *C. aerofaciens* into the 7β-BAs UCA and UDCA, respectively (Supp. Fig. 5). This exchange between species demonstrated multiple sequential transformations, where the products of one transformation became substrates for the next. Notably, neither 12-oxoBAs nor 3-oxoBAs were detected in coculture. In this and the previous coculture, the rapid BSH activity of *B. angulatum* and *C. aerofaciens* resulted in an accumulation of unconjugated BAs (CA, CDCA, and DCA), which were subsequently transformed into 7-oxoBAs by *B. theta*’s 7ɑ-HSD (Fig. 5C).

*S. infantarius* displayed very fast-acting BSH activity in both monoculture and coculture, deconjugating all conjugated BAs within six hours and generating unconjugated BAs that initially spiked and steadily decreased throughout the time course (Fig. 5). However, the anticipated production of 7-oxoBA in the *S. infantarius*-*B. thetaiotaomicron* coculture was limited. This observation could be explained by a loss of BAs to the *S. infantarius* bacterial membrane (Marion et al., 2018), a BA transformation not identified by our analysis, or by some other catabolic activity. In both monoculture and coculture, *S. infantarius* exhibited low-level transient accumulation of the MCBAs CDCA/DCA-Glutamine and CDCA-Glutamate between 3-6 hrs of growth. While BSHs were identified as the enzymes responsible for MCBA production (D. V. Guzior et al., 2024; Rimal et al., 2024), our observation of transient MCBA production with *S. infantarius* suggests that BSHs are also responsible for deconjugation of MCBAs, but further studies will be needed to confirm this possibility.

Surprisingly, when *B. thetaiotaomicron* was grown in pure culture, it accumulated a large amount, over 60 µM, of G-7-oxoLCA from GCDCA after 24 hrs, leaving all other conjugated BAs intact (Fig. 5B). When *C. symbiosum* was grown alone, it selectively deconjugated taurine-conjugated BAs after 24 hrs to release unconjugated BAs (Fig. 5A). In coculture, the slow-acting BSH activity of *C. symbiosum* allowed for *B. thetaiotaomicron* 7ɑ-HSD activity to transform conjugated BAs into glycine- and taurine-conjugated secondary BAs (Fig. 5C, Supp. Fig. 5). Conjugated secondary BA levels decreased after 24 hrs, indicating that they were subsequently deconjugated by *C. symbiosum* BSH to release 7-oxoBAs (Fig. 5C). If BSH activity had preceded HSD activity, we would have expected an accumulation of the unconjugated BAs CA, CDCA, and DCA before the production of the secondary 7-oxoBAs. Based on all time-series data, we concluded that HSD activity is more versatile than previously recognized, acting on both conjugated and unconjugated BAs to produce -oxoBAs. Ultimately, the composition of the resulting BA pool was determined by the combined activity and timing of BSH and HSDs from each bacterium.

### *B. thetaiotaomicron* HSD acts directly on conjugated primary BAs

As previously outlined, there are two potential pathways for the production of conjugated secondary BAs: 1) deconjugation by bile salt hydrolase (BSH), followed by secondary transformation by hydroxysteroid dehydrogenase (HSD) and subsequent reconjugation by BSH, or 2) direct transformation of conjugated BAs by HSD. To determine which pathway was responsible for the production of conjugated secondary BAs, we deleted the 7ɑ-HSD gene (Δ*hsd*) in *B. thetaiotaomicron* (Sherrod & Hylemon, 1977) using allelic exchange (García-Bayona & Comstock, 2019). If the multi-step pathway was occurring, knocking out the 7ɑ-HSD gene (Δ*hsd*) would result in an accumulation of CDCA due to deconjugation by BSH. However, if *B. thetaiotaomicron* 7ɑ-HSD acted directly on conjugated BAs, the provided conjugated BAs levels would remain unchanged.

We cultured both WT *B. thetaiotaomicron* and the Δ*hsd* mutant in monoculture and coculture with *C. symbiosum*, providing five conjugated BAs at 100 µM each (Fig. 6). In monocultures, the WT *B. thetaiotaomicron* strain transformed GCDCA to G-7-oxoLCA (Fig. 6B), while the Δ*hsd* strain did not perform any transformations, leaving all conjugated BAs intact by 72 hours (Fig. 6C). These results indicated that the 7ɑ-HSD in *B. thetaiotaomicron* directly dehydrogenated the C-7 hydroxyl group of the conjugated primary BA, GCDCA, producing the conjugated secondary BA, G-7-oxoLCA. This finding reveals a novel BA transformation route, challenging the established view that BAs must be first deconjugated before transformations on the BA core can occur (Supp. Fig. 1).

**Fig 6.**
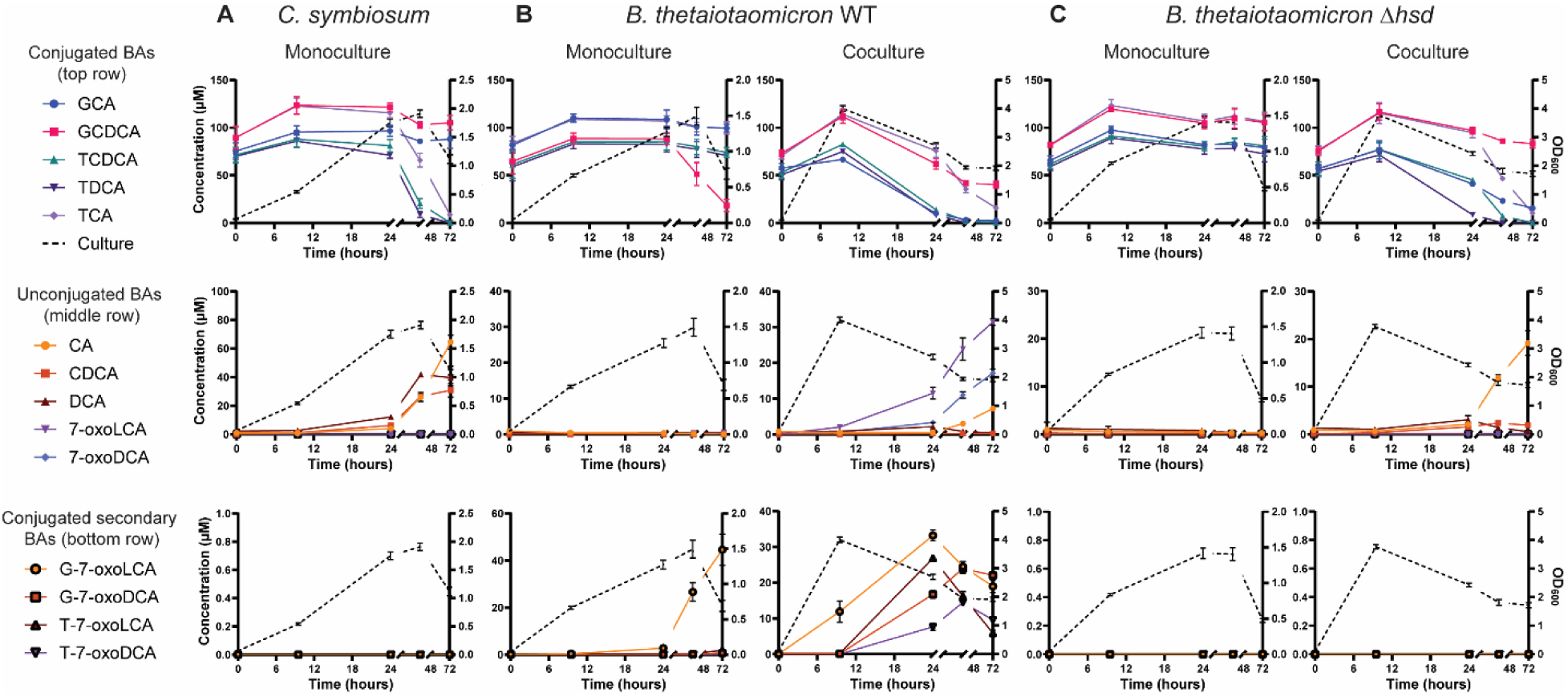
*B. thetaiotaomicron* 7ɑ-HSD makes conjugated secondary BAs. A) Bile acid transforming activity for *C. symbiosum* in monoculture. B) Bile acid transforming activity for WT *thetaiotaomicron* and *C. symbiosum* coculture. C) Bile acid transforming activity for *B. thetaiotaomicron* Δ*hsd* and *C. symbiosum* coculture. Species were cultured in triplicate and sampled over the course of 72 hours. Error bars represent the standard deviation of each averaged measurement. BA type is listed in the legend adjacent to each graph. Culture growth as measured by optical density is indicated by a black dashed line.

The coculture of *C. symbiosum* with WT *B. thetaiotaomicron* produced CA, DCA, 7-oxoLCA, 7-oxoDCA, and all four types of conjugated 7-oxoBAs (Fig. 6B). Given the diversity and abundance of conjugated secondary BAs, we tested this coculture using the *B. thetaiotaomicron* Δ*hsd* strain. The absence of *B*. *thetaiotaomicron* 7ɑ-HSD in the coculture resulted in an accumulation of CA and a lack of conjugated secondary BAs, again demonstrating the ability of its HSD to directly transform conjugated primary BAs (Fig. 6C). Since conjugated secondary BAs were absent, we confirmed that their production in the WT coculture was due to activity by *B*. *thetaiotaomicron* 7ɑ-HSD; however, *C. symbiosum* increased the diversity of conjugated secondary BAs. It remains unclear why *B*. *thetaiotaomicron* in coculture with *C. symbiosum* was able to produce several types of conjugated secondary BAs, when it only produced one in pure culture. Altogether, these data show that the 7ɑ-HSD of *B*. *thetaiotaomicron* is responsible for the production of conjugated secondary BAs in both monoculture from GCDCA and in coculture from GCDCA, TCDCA, GCA and TCA (Fig. 6).

## Discussion

Our investigation of bile salt hydrolase (BSH) and hydroxysteroid dehydrogenase (HSD) activity, two key bile acid (BA) transformations in phylogenetically diverse human gut bacteria, reveals that BSH activity is highly prevalent, with over 70% of the 77 tested strains exhibiting activity. Of these, 60% demonstrate varying substrate specificity for taurine- or glycine-conjugated BAs. Using coculture experiments, we demonstrate sequential transformations between bacterial species. highlighting the interplay between BSH and HSD activities in shaping BA pool diversity. We find that rapid and complete BSH activity, followed by HSD activity, drives the production of unconjugated secondary BAs. Conversely, limited secondary BA transformation activity leads to the accumulation of MCBAs. Unexpectedly, delayed or incomplete deconjugation activity allows HSDs to act on conjugated BAs, producing conjugated secondary BAs, representing a previously uncharacterized transformation (Supp. Fig. 1). These conjugated secondary BAs can subsequently undergo deconjugation by BSH, further diversifying the BA pool. Our findings challenge the conventional view of BSH activity as the single gateway reaction preceding other BA transformations, instead revealing its nuanced role in BA metabolism.

We further demonstrate that BSHs exhibit diverse dynamics, specificity, and sensitivity, broadening our understanding of the activity of this enzyme class. Historically, BSH activity has been associated primarily with the exponential growth phase, with a few exceptions noted in *Bacteroides* species (Begley et al., 2005; Ridlon et al., 2006). However, our time-series analysis shows that over one-quarter of the tested species decouple BSH activity from growth. Notably, we observe stationary-phase BSH expression in *C. symbiosum* (Fig. 5), a phenomenon not previously reported in *Clostridium* species. This observation indicates that while some BSHs are active during exponential growth or in response to BA exposure, others are active during the stationary phase, likely responding to alternative environmental cues.

In general, bacteria more completely deconjugate BAs at 100 µM compared to 500 µM, although exceptions, such as *L. scindens* and *B. dentium*, exhibit higher deconjugation efficiency at 500 µM (Fig. 3). This response may reflect a detoxification mechanism triggered by higher BA concentrations in these two species. Bacteria also exhibit stronger substrate preferences at elevated BA concentrations, possibly due to inhibited growth at high BA levels, as observed with *T. sanguinis* (Kemis et al., 2019). However, previous research suggests that BSH activity and reduced bacterial growth due to BA toxicity are unrelated properties, at least in the *Lactobacilli* (Moser & Savage, 2001). Overall, these findings suggest that enzyme capacity plays a role in BA deconjugation at physiologically relevant concentrations encountered in the gut.

In some species, such as *C. intestinalis*, *H. filiformis*, *D. formicigenerans*, and *E. ramosum*, secondary BA production by HSDs is limited despite robust BSH activity and the availability of unconjugated BA substrates. Similarly, *B. intestinalis*, *B. ovatus*, and *B. xylanisolvens* exhibit reduced HSD activity at 500 µM compared to 100 µM. The underlying causes of these differences in secondary BA production remain unclear but may result from regulatory mechanisms influenced by media composition or BA concentrations.

Our findings reveal that *B. thetaiotaomicron* oxidizes GCDCA to G-7-oxoLCA in pure culture when provided with conjugated BAs. This ability to produce conjugated secondary BAs is not restricted to *B. thetaiotaomicron*, as other members of the Bacteroidota, as well as *F. varium* (Fusobacteriodota), *C. intestinalis* (Actinomycetota), and *F. plautii* (Bacillota), appear to exhibit similar activity. Notably, *Bacteroides caccae*, *B. thetaiotaomicron* VPI-5482, and *B. thetaiotaomicron* 3731 produce conjugated secondary BAs more extensively at 500 µM BA concentrations than at 100 µM (Fig. 1). For these strains, increased HSD activity may play a role in detoxifying higher conjugated BA concentrations. For other species, however, the factors driving HSD activity, whether related to redox balance (Doden & Ridlon, 2021), detoxification (McMillan et al., 2023), or a combination of both, remain unclear.

Our findings on the production of conjugated secondary BAs align with prior studies showing that cell crude extracts or partially purified HSD enzymes from *Bacteroides fragilis* (Hylemon & Sherrod, 1975), *B. thetaiotaomicron* (McMillan et al., 2023; Sherrod & Hylemon, 1977), and *Clostridium limosum* (Sutherland & Williams, 1985) have HSD activity on both conjugated and unconjugated BAs. By integrating whole-cell assays, time-dependent LC-MS/MS-based BA measurements, and molecular genetics, our study expands upon these findings by directly validating HSD activity on conjugated BAs, highlighting the widespread promiscuity of HSDs and the potential relevance of conjugated secondary BAs *in vivo*.

Coculture experiments reveal that pairing two species can enable sequential, or additive BA transformations, as observed in cocultures of *B. thetaiotaomicron* with *B. angulatum, S. infantarius,* and *C. aerofaciens* (Fig. 5). Notably, in *B. thetaiotaomicron* and *C. aerofaciens* cocultures, the combination of 7ɑ-HSD and 7β-HSD activity leads to the production of urso-BAs. The factors governing HSD activity directionality remain poorly understood. However, our results align with previous findings that *C. aerofaciens* 7β-HSD switches from reduction to oxidation at ∼12 hrs of growth (MacDonald et al., 1982). Such reversible HSD activity may play a pivotal role in shaping the BA pool by redirecting BAs toward or away from other transformations. For example, the dehydroxylation of BAs to produce DCA and LCA cannot occur on -oxoBAs. These urso-BAs, which are more hydrophilic and less toxic to both the microbiota and the host due to their hydrophilicity (Watanabe et al., 2017), have therapeutic relevance for biliary disorders (Tang et al., 2018), heart disease (Hanafi et al., 2018) and various cancers (Goossens & Bailly, 2019).

When HSD activity is limited and BSH activity is high, MCBAs are often produced by the enzyme’s recently described BSH acyltransferase activity (D. V. Guzior et al., 2024; Rimal et al., 2024). Consistent with prior studies, higher concentrations of MCBAs correlates with greater diversity (D. V. Guzior et al., 2024). Certain species, such as *Ruminoccocus gnavus*, *B. bifidum*, and *E. bolteae* produce robust and diverse MCBAs, whereas others like *L. scindens*, *Holdemanella hathewayi*, and *C. symbiosum* show more specificity and produce fewer MCBAs (Daly et al., 2021; D. V. Guzior et al., 2024; D. V. Guzior & Quinn, 2021; Lucas et al., 2021). In general, more species were capable of producing MCBAs from conjugated BAs than from unconjugated BAs based on our prior study (Lucas et al., 2021). Increased production of MCBAs may be due to greater induction of BSH by the presence of conjugated BAs.

Similar to how BSHs exhibit specificity for glycine- or taurine-conjugated BAs, they also demonstrate specificity in the types of MCBAs they produce. Species with BSH specificity for glycine or taurine may exhibit deconjugating activity yet not produce MCBAs with alternative amino acid conjugations in our study. MCBA conjugation profiles may not only be shaped by BSH specificity but also by autotrophic amino acid production of each species (Rimal et al., 2024). While glycine-conjugated MCBAs are known to be prevalent, their production could not be measured in this study because glycine-conjugated BAs (GCA and GCDCA) were substrates in the media.

The relationship between bioinformatically predicted BSH activity and actual BA deconjugation is complex and often inconsistent. Several species exhibit deconjugating activity even though no identifiable BSH homologs are found, suggesting that non-homologous enzymes may be responsible for BA deconjugation in these organisms. This observation is noted for *H. filiformis, C. intestinalis, E. ramosum, C. symbiosum, D. formicigenerans, B. hansenii, B. stercoris, B. caccae, B. cellulosilyticus,* and *B. thetaiotaomicron* 3731. Interestingly, all these species, except for *D. formicigenerans*, preferentially deconjugate taurine-conjugated BAs, providing insights into the evolutionary lineage and specialization of taurine-specific BSHs. In contrast, *P. rettgeri, P. penneri*, and *P. mirabilis* of the Pseudomonadota, and *C. sporogenes* and *B. hydrogenotrophica* of the Bacillota, possess bioinformatically identified BSHs but do not display measurable activity. This lack of activity may result from the absence of functional transporters for importing conjugated BAs, the presence of genetic alterations that render the enzyme inactive, or the lack of BSH expression under the environmental conditions tested in our study. These findings highlight that while bioinformatic predictions are highly informative, they are not sufficient to fully identify BA transforming activity, particularly when novel enzymes, regulatory differences, or uncharacterized transport systems may be involved.

Bacterial BA transformations have long been recognized for their role in converting a limited set of host-synthesized BAs into hundreds, if not thousands, of modified derivatives (Mohanty et al., 2024). These diverse BAs vary in their capacity to facilitate fat absorption, shape the gut microbial community, and interact with host receptors. Traditionally, BA transformation has been described as a linear process in which primary conjugated BAs are first deconjugated, releasing free primary BAs that can subsequently be modified into secondary BAs by gut bacteria. However, the recent discovery that gut bacteria can conjugate BAs to glycine, reforming conjugated “primary” BAs through microbial action, shifts our understanding of microbial contributions to the BA pool and expands the known activity of BSH.

In this study, the identification of conjugated secondary BAs reveals a novel BA transformation and challenges the long-held assumption that deconjugation is a prerequisite for further transformation. Our findings continue to overturn conventional views of the BA transformation process and underscore the need to move beyond the concept of a linear “pathway.” Instead, we propose a transformation “network” model that accounts for the timing, specificity, and interconnected nature of BA modifications within a dynamic and ever-changing BA pool. Such a framework will provide a more accurate and relevant understanding of bacterial activity in *in vivo* systems.

Systematic investigations of *in vitro* BA transformations bring us closer to understanding and interpreting *in vivo* BA pools associated with metabolic diseases, gastrointestinal cancers, and improved health outcomes post-bariatric surgery. Our findings highlight the remarkable diversity of BAs and the transformations that produce them, emphasizing their potential for manipulation to improve human health. However, the observed variation in BA deconjugation and transformation, even within species of the same genus, underscores the limitations of making generalized statements about BA toxicity, growth effects, and microbial activity. In addition, environmental factors such as media composition, pH, and microbial interactions contribute to discrepancies between studies and limit the applicability of *in vitro* observations to *in vivo* systems.

To address these challenges, further studies are needed to identify the mechanisms that regulate BA-transforming activity in gut bacteria, including the roles of BA transporters, BSH expression levels, and cell death in shaping the dynamic BA network. Systematic, time-course analyses using diverse BA substrates under physiologically relevant conditions are essential to unravel the regulatory pathways and environmental cues that drive these activities. A deeper understanding of these factors is critical if we are to reliably manipulate the BA pool to promote beneficial health outcomes.

## Materials and Methods

### Strains

All strains are listed in Supp. Table 1. Most strains are previously sequenced and come from the Human Microbiome Project. Further information for strains isolated in our lab: Strain “1RE7” was isolated from an anaerobic enrichment in a medium supplemented with rutin, inoculated with a human fecal sample. The strain consumes both rutin and quercetin. The sequence of the full-length 16S gene is 96% identical to that of *C. scindens* CG19-1. Strain “J02” was isolated from an anaerobic enrichment in medium supplemented with rutin, inoculated with a human fecal sample (WLS #82). The sequence of the 16S gene is >99% identical to that of *E. tayi* strain B086562 (783/784 bases match). Strain “K01” was isolated from an anaerobic enrichment in media supplemented with rutin, inoculated with a human fecal sample. The sequence of the 16S gene is >99% identical to that of *Enterococcus durans* JCM8725 (900/901 bases match) and similarly matches many *E. faecium* strains (902/903 bases match). A full-length 16S gene sequence might be more definitive. Strain “J01” was isolated from an anaerobic enrichment in media supplemented with quercetin, inoculated with a human fecal sample. The sequence of the 16S gene is >99% identical to that of several *Enterococcus* species (*lactis*, *durans*, *faecium*) all with 870/871 bases matching. A full-length 16S gene sequence might be more definitive. Strain “K02” was isolated from an anaerobic enrichment in medium supplemented with rutin, inoculated with a human fecal sample (WLS #10). The sequence of the 16S gene is >100% identical to multiple *P. mirabilis* strains (823/823 bases match). Strain “L02” was isolated from an anaerobic enrichment in media supplemented with quercetin, inoculated with a human fecal sample. The sequence of the 16S gene is >99% identical to that of several *S. anginosis* strains (854/856 bases match).

### Media

For the systematic BSH analysis, all strains were grown on Colossal Mega Medium, which was filter-sterilized and stored in a Coy anaerobic chamber (5% H_2_, 20% CO_2_, and 75% N_2_) at least 24 hours prior to use. Colossal Mega Medium contains (per liter tap distilled water): 100 mL (1M, pH 7.2) potassium phosphate buffer, 10 g tryptone peptone, 5 g yeast extract, 5 g meat extract, 4 mL (25 mg/100 mL) Resazurin, 1.8 g D-glucose, 0.9 g D-maltose, 0.86 g D-cellobiose, 0.46 g D-fructose, 2 g sodium acetate trihydrate, 0.02 g MgSO_4_ꞏ7 H_2_O, 2.1 g NaHCO_3_, 0.08 g NaCl, 1 mL (0.8g/100mL) CaCl_2_, 1 mL (1 mg/mL in 100% ethanol) vitamin K_3_ (menadione), 1 mL (1.2mg hematin/mL in .2M histidine, pH 8.0) histidine hematin, 2 mL (25% vol/vol) Tween 80, 10 mL ATCC MD-VS vitamin mix, 10 mL ATCC MD-TMS trace mineral mix, 1 mL (40mg/100mL) FeSO_4_ꞏ7 H_2_O, and 0.5 g L-cysteineꞏHCL. This specific medium was designed to allow growth for all species in this study. Additions and modifications for specific strains were as follows: For cultures of *Akkermansia muciniphila* the medium was amended with 1 mg/mL mucin. For cultures of *Clostridium orbiscindens* the medium was amended with lysine.

For time-course cocultures, all strains were grown on Low Yeast Extract (LYE) Medium, which was made anaerobic using a triple-vacuumed pressure bottle before being brought into a Coy anaerobic chamber (5% H_2_, 20% CO_2_, and 75% N_2_), and then filter-sterilized. Low Yeast medium contains (500 mL Milli-Q water): 50 mL (1 M, pH 7.0) potassium phosphate buffer, 0.36 g tricine, 2.0 mL (0.025%) resazurin, 1 g yeast extract, 0.5 mL (25% [vol/vol]) tween 80, 3.4 g sodium acetate trihydrate (FW 136), 0.55 g sodium succinate hexahydrate (FW 270), 1.46 g sodium chloride (FW 58.44), 0.54 g ammonium Chloride (FW 53.49), 3.6 g d-glucose (FW 180.16), 1.8 g d-maltose (FW 360.3), 1.0 mL (0.5 M) potassium sulfate, 1.0 mL (1.0 M) magnesium chloride hexahydrate (MgCl_2_. 6H_2_O), 0.2 mL (1.0M) calcium chloride dihydrate (CaCl_2_.2H_2_O), 1.68 g sodium bicarbonate (FW 84.0), 0.5 mL (1.2 mg hematin/ml in 0.2 M histidine, pH 8.0) histidine hematin solution, 0.125ml vitamin K1+ K3 solution (used “2x” stock), 10 mL ATCC MD-VS vitamin mix, 5 mL 50x trace mineral mix solution [0.29 mL (30 µM) MnCl2.4H_2_O, 0.06 mL (10 µM) ZnCl_2_, 0.047 mL (4 µM) CoCl2.6H_2_O, 0.012 mL (1 µM) Na_2_MoO4.2H_2_O, 0.008 mL (1 µM) Na_2_SeO_3_, 0.059 mL (5 µM) NiCl_2_.6H_2_O, 0.016 mL (1 µM) Na_2_WO_4_.2H_2_O, adjust volume to 1 L, store under N_2_, refrigerated], 1 mL ferrous sulfate heptahydrate (FeSO4ꞏ7H2O), and 0.25 g l-cysteine HCl. Adjust pH to ⁓7.3 -7.1.

### Sample handling and growth conditions

For the systematic investigation reported in Figures 1, 2, 3 and Supplementary Figure 3, cultures were started from freezer stocks and grown overnight to a high density (O.D. 600 range of .349-1.9, measured directly in the tube) in Hungate tubes containing Colossal Mega Medium with an atmosphere of 75% N_2_, 20% CO_2,_ 5% H_2_ at 37°C. These cultures were then used to inoculate (1:15 dilution) 3 mL of Colossal Mega Medium in Hungate tubes amended with bile acids. There were 2 sets of conditions; media contained 100 μM or 500 μM of each of the five conjugated bile acids, glycocholic acid (GCA), glycochenodeoxycholic acid (GCDCA), taurocholic acid (TCA), taurochenodeoxycholic acid (TCDCA), and deoxycholic acid (DCA). Each condition was tested in duplicate. In addition, we tested for spontaneous bile acid degradation or transformation in uninoculated controls containing bile acids. Once cultures reached stationary phase, 1 mL of culture was collected, spun down at room temperature for 10 minutes at 10,000 g, and the supernatant transferred to a fresh tube. Using HPLC-grade H_2_O, the supernatants were diluted 1:200 or 1:1000 for the 100 μM or 500 μM conditions, respectively. After dilution, 100 μL were transferred to an HPLC vial for analysis.

For the monoculture and coculture time course analyses, individual freezer stocks were inoculated into Colossal Mega Medium and grown overnight to a high bacterial density (OD_600_ > 2). The following day, multiple dilutions were made for each culture in LYE medium containing 0.1% yeast extract and allowed to grow overnight. Cultures in exponential phase were then used as inoculum. Growth curves and bile acid measurements for Figure 4 and Supplementary Figure 4 were performed in 125 mL Erlenmeyer flasks containing 30 mL of 0.1% LYE medium. Growth curves and bile acid measurements for Figures 5 and 6 were performed in Hungate tubes containing 10 mL of 0.1% LYE medium. Both sets were amended with 100 µM each of GCA, GCDCA, TCA, TCDCA, and TDCA. Monoculture starting ODs were 0.05 and cocultures were started with an equal proportion of both monocultures (total OD being ∼ 0.1). For *B. thetaiotaomicron* and *C. symbiosum*, monocultures starting ODs were 0.03 and 0.12, respectively, with the coculture OD at ∼ 0.15. Uninoculated controls containing bile acids were used to assess spontaneous bile acid degradation and transformation. All experiments were performed in triplicate. Samples were drawn at multiple timepoints including “zero” time point based on the growth pattern: rigorous sampling was done during exponential phase (7-8 timepoints) and 4-5 timepoints were included in stationary phase. 0.3 mL of culture was collected at each timepoint and spun down, and the supernatant was diluted at 1:100 using HPLC-grade H_2_O for analysis by HPLC-MS.

### uHPLC-MS/MS measurements

Samples were analyzed using an ultra-high pressure liquid chromatography-tandem mass spectrometry (uHPLC-MS/MS) system consisting of a ThermoScientific Vanquish uHPLC system coupled to a heated electrospray ionization (HESI; using negative polarity) and hybrid quadrupole high resolution mass spectrometer (Q Exactive Orbitrap; Thermo Scientific). Settings for the ion source were: auxiliary gas flow rate of 10, sheath gas flow rate of 30, sweep gas flow rate of 1, 2.5 kV spray voltage, 320°C capillary temperature, 300°C heater temperature, and S-lens RF level of 50. Nitrogen was used as nebulizing gas by the ion trap source. Liquid chromatography (LC) separation was achieved using a Waters Acquity UPLC BEH C18 column with 1.7 μm particle size, 2.1 × 100 mm in length. Solvent A was water with 10 mM ammonium acetate adjusted to pH 6.0 with acetic acid. Solvent B was 100% methanol. The total run time was 31.5 min with the following gradient: a 0 to 24 min gradient from 30% solvent B (initial condition) to 100% solvent B; held 5 min at 100% solvent B; dropped to 30% solvent B for 2.5 min re-equilibration to initial condition. The flow rate was 200 μL/min throughout. Other LC parameters were as follows: autosampler temperature, 4°C; injection volume, 10 μL; column temperature 50°C. The MS method performed a full MS1 full-scan (290 to 1000 m/z) together with a series of PRM (parallel reaction monitoring) scans.

Identity of conjugated secondary BAs was confirmed through LC-MS/MS fragmentation. The MS method performed a full MS1 full scan (290 to 2,000 *m/z*) together with a series of parallel reaction monitoring (PRM) scans in positive mode. These MS2 scans (selected-ion fragmentation) were centered at *m/z* values of 448, 464, 498, and 514. Fragmentations were performed at 30 normalized collision energy (NCE). All scans used a resolution value of 17,500, an automatic gain control (ACG) target value of 1E6, and a maximum injection time (IT) of 40 ms. Experimental MS data were converted to the mzXML format and used for bile acid identification. Bile acid peaks were identified using MAVEN (metabolomics analysis and visualization engine) (Clasquin et al., 2012; Melamud et al., 2010).

### Determination of bile acid concentrations

Bile acid quantitation was achieved using standard concentrations of each bile acid ranging from .0625 to 2 μM to generate six-point external standard curves. The detection limit was below 0.01 μM for all bile acids. The threshold for reported core bile acid transformations was 0.008 μM. Standards were purchased from Avanti Polar Lipids and dissolved and stored in methanol at -80 °C. See Table S2 for bile acid standard names and structural features. For MCBAs, compounds were identified by their exact mass (mass error of less than 2 parts per million) and previously determined retention times. Values were presented as z-scores to demonstrate relative abundance. Conjugated secondary BA concentrations were estimated using the commercially available glyco-12-oxolithocholanic acid (G-12-oxoLCA) and tauro-12-oxolithocholanic acid (T-12-oxoLCA). For BA measurements in time series-analyses unconjugated BAs were normalized to conjugated BA measurements in uninoculated controls.

### *In silico* analysis

Access the genome sequence in NCBI for 77 bacterial strains. Get all CDS genes from the 77 available genomes (amino acid sequences). Use curated 84 BSH genes from Foley et al. (PMID: 36914755) as BSH reference database for BLASTp. Use the same 84 BSH genes to build the HMM (Hidden Markov Model) to reserve BSH conserved domains when predicting BSH genes. Use the following criteria to determine the BSH from all CDS: gene length between 300 to 400 bp; has at least one BLASTp to BSH reference genes (identity > 25%); has hit to BSH HMM (full sequence score > 100). Clustal Omega was used for multiple alignment of predicted BSH genes, taurine- or glycine-preferring BSH was predicted by a 3-residue selectivity loop: taurine-preferring BSH contain ‘G-X-G’ motif and glycine-preferring BSH contain ‘S-R-X’ motif.

### Generation of *B. thetaiotaomicron* VPI-5482 *hsd* mutant

An in-frame hydroxysteroid dehydrogenase (hsd) deletion mutant was generated using a previously described counter-selectable allelic exchange principle (García-Bayona & Comstock, 2019). Briefly, ∼1 kilobase upstream (including the start codon) and downstream (including the stop codon) fragments of *hsd* open-reading frame were amplified using the high fidelity Herculase polymerase and the primer pairs TAAGATTAGCATTATGAGTGGAAAAGAAAAAGTGATCTGG and ATATTTATGACATATATGTTGAGAATTTGATGATTAC; and CAACATATATGTCATAAATATACCCCGGAC and CGAATTCCTGCAGCCCGGGGATATAAGCGTACGAGGTG, respectively. These amplified fragments were cloned into the BamHI site of the suicide vector pLGB13 via Gibson assembly. The resulting construct was transformed into *E. coli S17-1 λ pir* strain. After cloning, the junction sequence was verified by Sanger sequencing.

This vector was introduced into *B. thetaiotaomicron* by biparental mating (conjugation) between *E. coli* and *B. thetaiotaomicron* and single-crossover events were selected on CMM-blood agar plates containing gentamicin (20 µg/mL) and erythromycin (10 µg/mL) to enrich exconjugants. Resulting colonies were purified twice on the same plates. The cultures from the purified colonies were further plated on plates containing gentamicin and anhydrotetracycline (100 ng/mL), a counter-selection marker. To identify isolates that had lost the gene, colonies derived from a single original colony were screened by PCR. About 25% colonies were devoid of *hsd* gene, and one such colony was used following the sequence and functional verification.

## Supporting information

Supplementary Table 1. Bacterial strain and bile acid information.

Supplementary Table 2. Raw data for Fig. 1, Fig. 2, and Supp. Fig 3.

## Acknowledgments

This work was supported in part by the National Institutes of Health (NIH) grants HL148577 (F.E.R.) and by the Transatlantic Networks of Excellence Award from the Leducq Foundation.

L.N.L. was supported by The Molecular and Applied Nutrition Training Program (MANTP) NIH T32 DK 007665.

L.E.C. was supported by the National Institute of General Medical Sciences of the National Institutes of Health under award numbers T32GM135066. L.E.C. was also supported by the University of Wisconsin—Madison SciMed Graduate Research Scholars Fellowship.

