## Supplementary Figures 1-5 for "Investigation of Bile Salt Hydrolase Activity in Human Gut Bacteria Reveals Production of Conjugated Secondary Bile Acids"

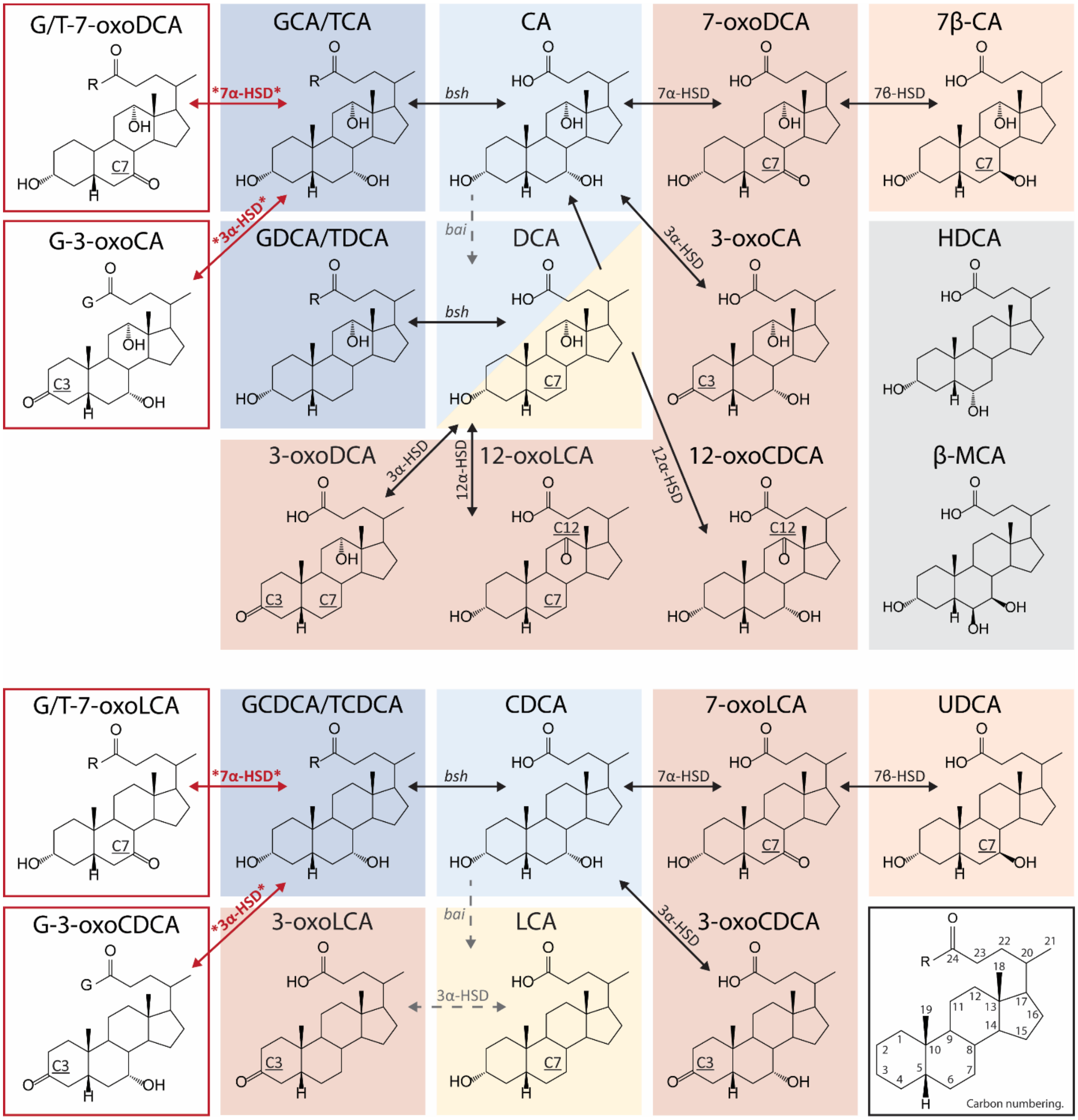


**Supplementary Figure 1. Bile acid transformations and structures.** Transformations by *bsh, bai,* and *hsd*. Observed transformations are indicated by a solid black arrow, novel transformations are indicated by a bold red arrow with asterisks, and known but unobserved transformations are indicated by a grey dashed arrow. Carbon atom numbering for bile acid steroid core is shown in a black box in the bottom right corner.

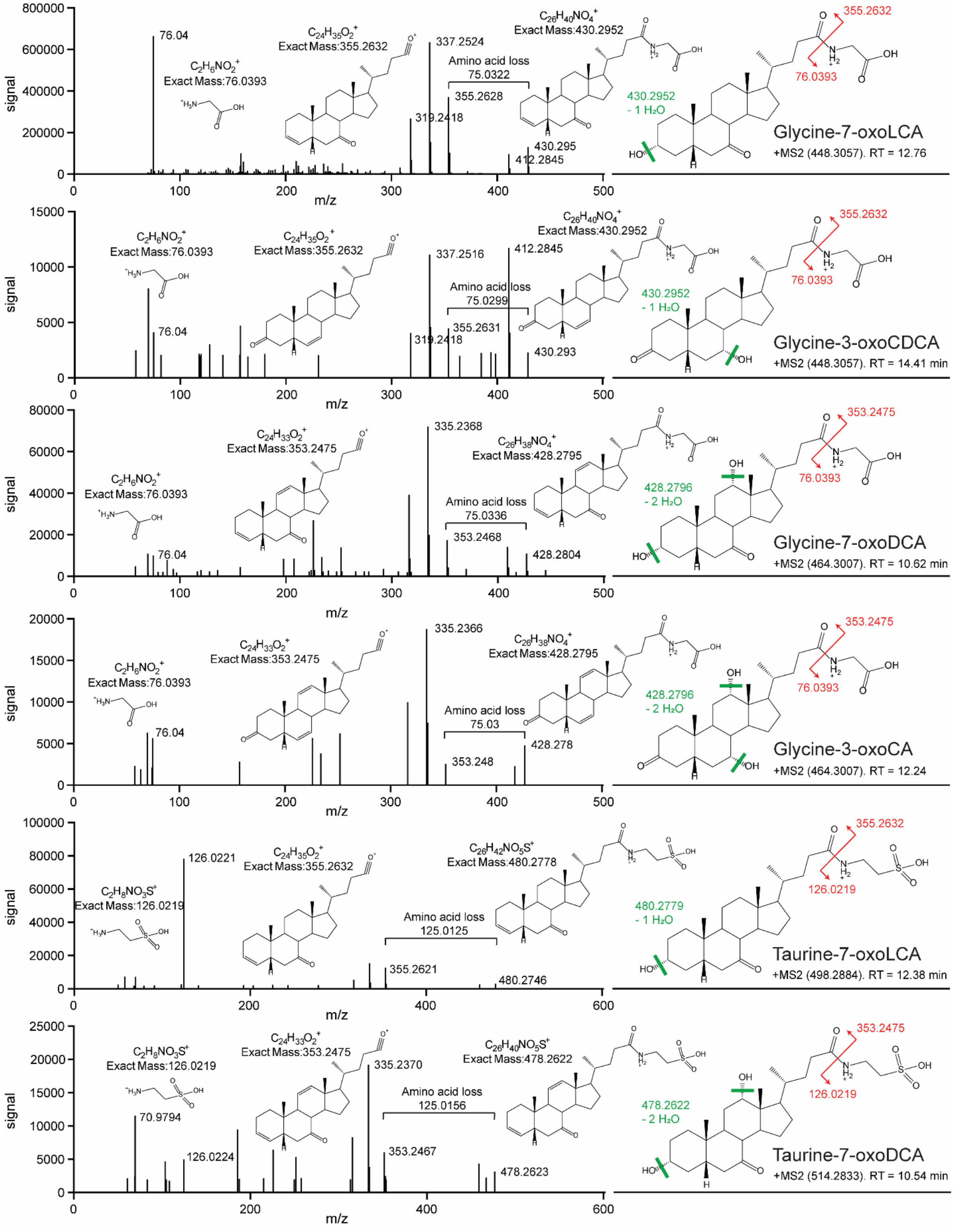


**Supplementary Figure 2. MS/MS spectra for observed conjugated secondary bile acids.** Parent ion structure, mass, and retention times are listed along with the structure and exact masses for three identifying fragments: the major sterol fragment, the fragment resulting from amino acid loss, and the amino acid fragment. See methods section called “uHPLC-MS/MS measurements”.

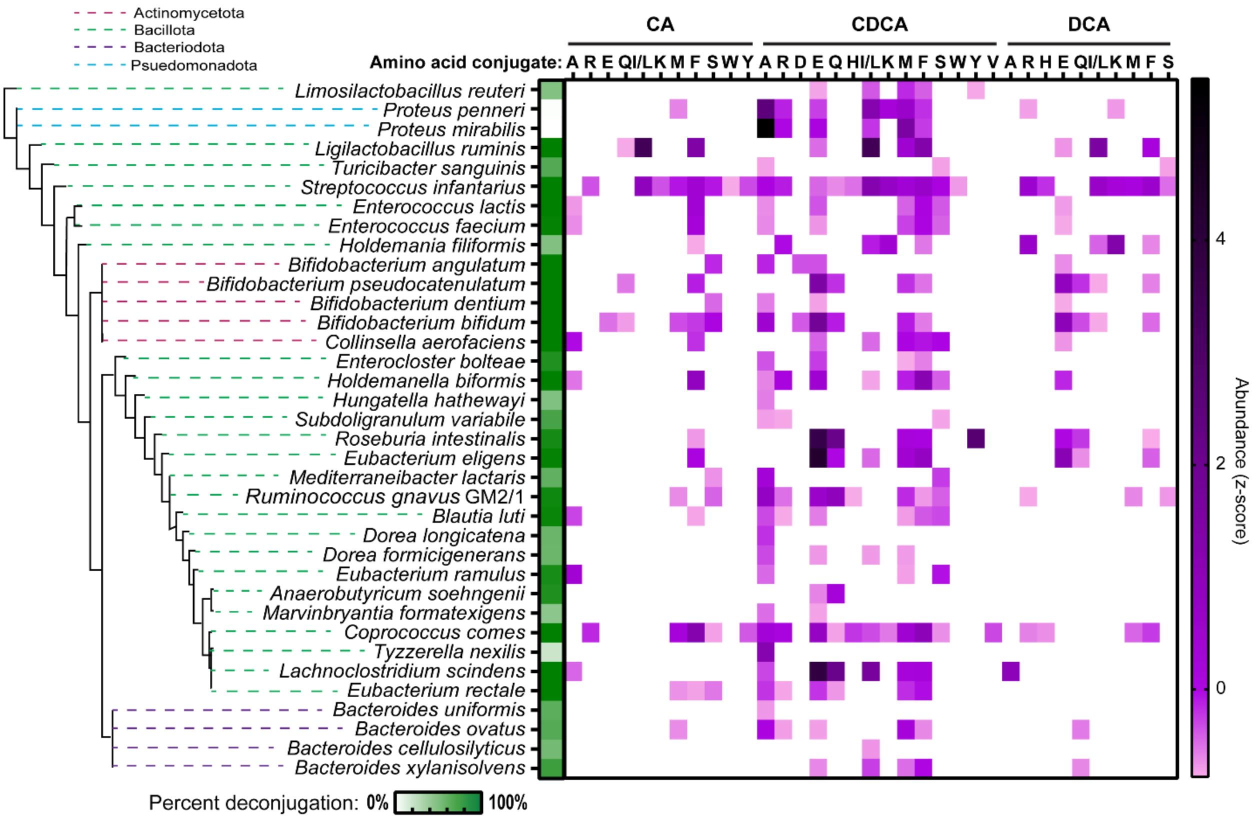


**Supplementary Figure 3. Microbially conjugated bile acid production at 500 µM.** The heat map shows relative levels of MCBAs produced, as denoted by z-score, with the mean and standard deviation calculated from raw signal across samples. Phyla information is indicated by color-coded dashed lines in the phylogenetic tree. Amino acid conjugations are indicated by their single letter abbreviation across the top of the heat map.

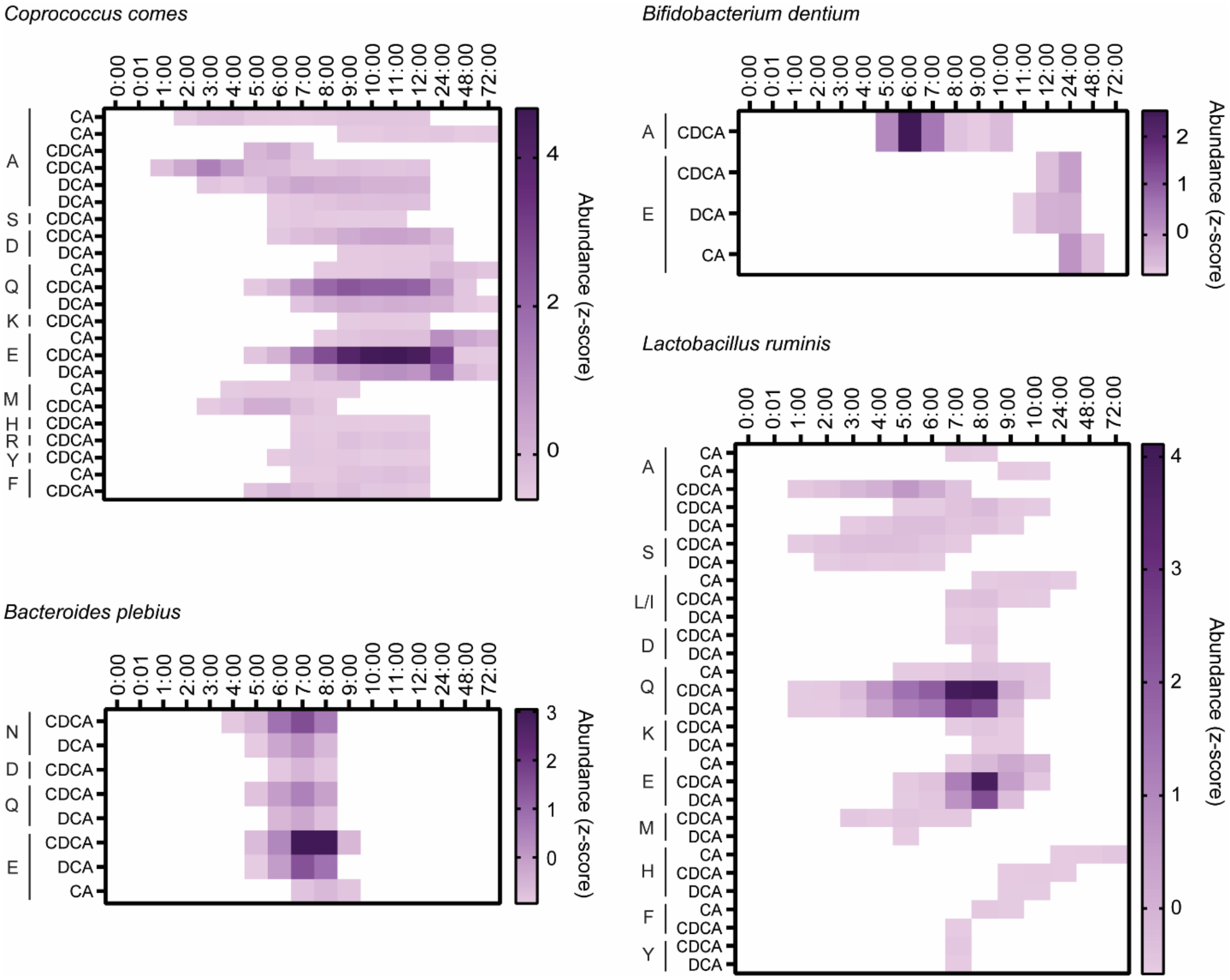


**Supplementary Figure 4. Microbially conjugated bile acid production for monocultures.** The heat map shows relative levels of MCBAs produced, as denoted by z-score, with the mean and standard deviation calculated from raw signal for each species. Amino acid conjugations are indicated by their standard single letter abbreviation to the left of each bile acid core.

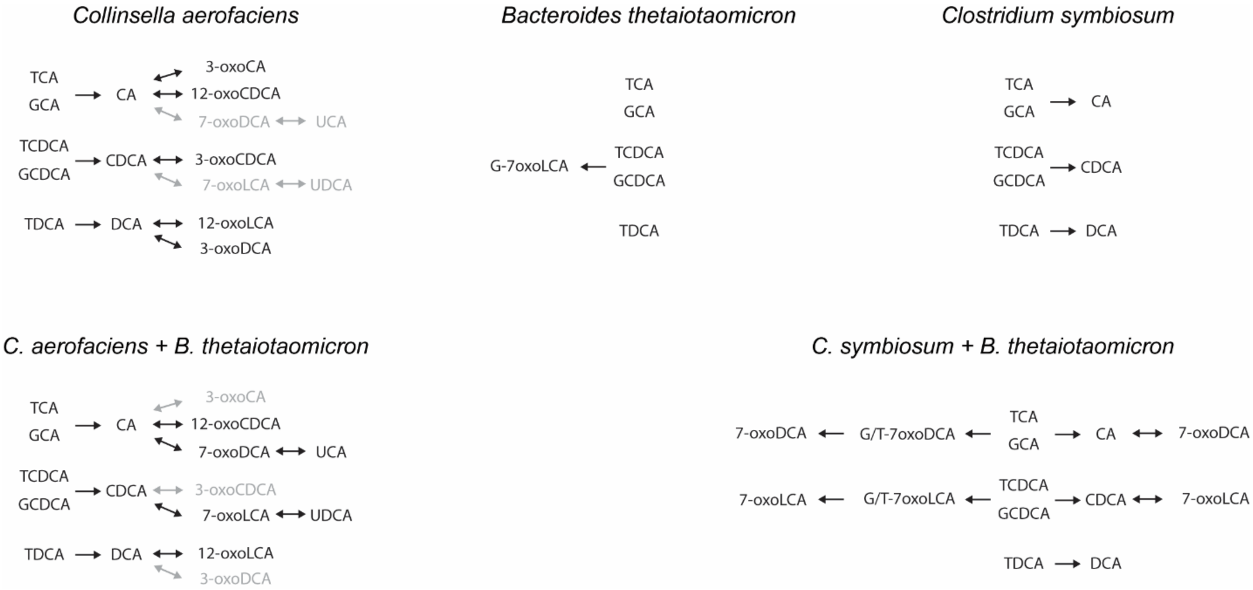


**Supplementary Figure 5. Bile acid transformation networks for cocultures of *B. thetaiotaomicron* with *C. aerofaciens* and *C. symbiosum.*** A visual aid to compare monoculture transformation networks with coculture transformation networks.
